# Uneven missing data skews phylogenomic relationships within the lories and lorikeets

**DOI:** 10.1101/398297

**Authors:** Brian Tilston Smith, William M. Mauck, Brett Benz, Michael J. Andersen

## Abstract

The resolution of the Tree of Life has accelerated with advances in DNA sequencing technology. To achieve dense sampling, it is often necessary to obtain DNA from historical museum specimens to supplement modern genetic samples. However, DNA from historical material is generally degraded and fragmented, which presents various challenges. In this study, we evaluated how the coverage at variant sites and missing data among historical and modern sample types impacts phylogenomic inference. We explored these patterns in the brush-tongued parrots (lories and lorikeets) of Australasia by sampling ultraconserved elements in 105 taxa. Trees estimated with low coverage sites had several clades where historical or modern samples clustered together, which were not observed in trees with more stringent filtering. To assess if the aberrant relationships were affected by missing data, we performed a targeted outlier analysis of sites and loci and a more general data reduction approach where we excluded sites based on a percentage of data completeness. The outlier analyses showed that 6.6% of total sites were driving the topological differences among trees built with and without low coverage sites, and at these sites, historical samples had 7.5x more missing data than modern ones. An examination of subclades identified loci biased by missing data, and the exclusion of these loci shifted phylogenetic relationships. Predictive modeling found that outlier analysis scores were not correlated with summary statistics of locus alignments, indicating that outlier loci do not have characteristics differing from other loci. Excluding missing data by percentage completeness indicated that sites with 70% completeness were necessary to avoid spurious relationships, but more stringent conditions of data completeness produced less-resolved trees. After accounting for biased loci and understanding the stability of relationships, we inferred a more robust phylogenetic hypothesis for lories and lorikeets.

## Introduction

Historical and ancient DNA from museum specimens is widely employed for incorporating rare and extinct taxa into phylogenetic studies (e.g., Thomas et al. 1989; Mitchell et al. 2014; Fortes et al. 2016). The inclusion of these samples has helped discover and delimit species (Helgen et al. 2013; Paijmans et al. 2017), resolve phylogenetic relationships (Mitchell et al. 2016), and clarify biogeographic history (Kehlmaier et al. 2017; Yao et al. 2017). DNA sequences obtained from dry and alcohol-preserved museum specimens have been collected using a range of techniques, including Sanger sequencing (Sorensen et al. 1999), restriction site-associated DNA sequencing (Tin et al. 2015; Ewart et al. 2019), and sequence capture of reduced (McCormack et al. 2016; Ruane et al. 2017; Linck et al. 2017) or whole genomes (Hung et al. 2014; Enk et al. 2014). However, DNA sequences collected from these museum specimens are subject to errors associated with contamination (Malmström et al. 2005), DNA degradation (Briggs et al. 2007; Sawyer et al. 2012), and low coverage in read depth (Tin et al. 2015), which all present challenges in distinguishing evolutionary signal from noise.

Sequence capture of ultraconserved elements (UCEs) is a popular approach for collecting orthologous genomic markers in phylogenomic studies (Faircloth et al. 2015; Chakrabarty et al. 2017; Esselstyn et al. 2017) and is increasingly used for historical specimens (McCormack et al. 2016; Hosner et al. 2016; Ruane et al. 2017). A common finding is that the loci recovered are typically shorter in older samples (McCormack et al. 2016; Hosner et al. 2016; Ruane et al. 2017). Shorter loci are potentially problematic because the sequence capture approach targets the invariable UCE core, limiting the portion of the flanking region that contains polymorphic sites. Another factor that may cause differences among historical and modern samples is that phylogenomic pipelines that do not involve variant calling typically employ read-specific filtering, where the average read depth across all sites in a locus is used to determine whether the locus is excluded. Under this type of scenario, low coverage sites may pass typical filters, exacerbating differences among historical and modern samples. Although some studies only use DNA sequences collected from historical or ancient samples (e.g., Hung et al. 2014), most phylogenetic approaches involving non-contemporaneous samples combine with those from modern samples. For those that do use DNA from both sample types, additional challenges in downstream analyses may arise due to an asymmetry in the phylogenetic signal caused by non-random missing data (e.g., Hosner et al. 2016).

The impact of missing data on phylogenetic inference remains contentious (Lemmon et al. 2009; Simmons 2012; Wiens and Morrill 2011; Hovmöller et al. 2013; Simmons 2014; Jiang et al. 2014; Streicher et al. 2015). Missing data have been shown to bias phylogenetic relationships, particularly when the missing sites are non-randomly distributed (e. g., Lemmon et al. 2009; Simmons 2012; Simmons 2014) However, findings also suggest that even when some taxa have a large proportion of sites with no data, phylogenetic signal is retained if enough characters are present (Philippe et al. 2004; Roure et al. 2012; Shavit Grievink et al. 2013; Molloy and Warnow 2017). Bias may manifest as inflated support values and erroneous branch lengths, or as inconsistency between optimality criteria or phylogenomic approaches (i.e., concatenation vs the multi-species coalescent). The increased availability of phylogenomic data has provided a more nuanced look at missing data’s effect on phylogenetic inference (Philippe et al. 2004; Huang and Knowles 2014; Streicher et al. 2015; Xi et al. 2015). One means of dealing with missing data in phylogenomic datasets is to filter loci based on the proportion of either missing sites or missing species in the dataset (Hosner et al. 2016). However, this approach may not directly target problematic regions of an alignment, and phylogenetically informative signal may be discarded unnecessarily. A more direct approach would entail identifying which specific sites or genes are influenced by missing data.

Analyses of outlier sites or loci in phylogenomic data indicate that a small portion of genes can have a large impact on a topology (Shen et al. 2017; Arcila et al. 2017; Brown and Thomson 2017; Walker et al. 2018). These conflicting genealogies can be due to biological processes (e.g., incomplete lineage sorting; introgression; horizontal gene transfer) or to spurious phylogenetic signal caused by poor alignments, paralogy, and/or sequencing error. Putative outlier loci have been identified using topology tests (Arcila et al. 2017; Esselstyn et al. 2017), Bayes factors (Brown and Thomson 2017), and site/gene-wise log-likelihood differences among alternative topologies (Shen et al. 2017; Walker et al. 2018). Support for particular phylogenetic hypotheses may be driven by a small subset of loci (Brown and Thomson 2017; Walker et al. 2018), and the targeted removal of outlier loci can reconcile differences among topologies. Outlier analyses provide a framework for assessing how differences between historical and modern DNA sequences impact phylogenetic inference. In this study, we performed site and gene likelihood outlier analyses to evaluate whether sequence coverage and missing data impact phylogenetic relationships in our focal group, the Loriini.

Lories and lorikeets, commonly known as the brush-tongued parrots, are a speciose clade (Tribe: Loriini) of colorful birds that are widely distributed across the Australasian region (Forshaw et al. 1989). The characterization of distributional ranges, phenotypic variation, and systematics of the clade was the product of expansive biological inventories that peaked during the early 1900s (Mivart 1896; Forshaw et al. 1989). The geographical extent of this work encompasses thousands of large and small islands spread across many countries in the Australasian region. Given these immense logistical constraints, modern collecting expeditions that aim to produce voucher specimens with genetic samples for continued systematic work (e.g., Kratter et al. 2006; Andersen et al. 2017) have been much more focused in scope relative to the pioneering work of the 20th century that produced extensive series of specimens across species’ entire ranges (e.g., Mayr 1933; Mayr 1938; Mayr 1942; Amadon 1943). Thus, the lack of modern genetic samples means that phylogenetic relationships in many groups, like the Loriini, remain unresolved. To get around this constraint, phylogenomic studies have sourced DNA from historical specimens to fill modern sampling gaps (Moyle et al. 2016; Andersen et al. 2018).

Prior phylogenetic work on the Loriini showed evidence for at least three paraphyletic genera (*Trichoglossus*, *Psitteuteles*, and *Charmosyna*) and highlighted the need for increased taxon and genomic sampling to more fully resolve relationships among taxa (Schweizer et al. 2015). To this end, we collected UCEs from 105 described taxa in the Loriini. Our sampling design used DNA isolated from fresh tissues (hereafter modern) and historical specimens, including some over 100 years old (hereafter historical; supplementary fig. S1; supplementary table S1). We anticipated challenges with processing, recovering, and analyzing UCEs from historical specimens, and expected that biases in the DNA sequence data might yield misleading relationships. To evaluate biased phylogenetic signal and explore options for maximizing the amount of data recovered, we produced alignments using different site coverage thresholds that produced alignments with varying levels of missing data. We then estimated phylogenies with particular sites and loci removed, and with varying percentages of data completeness to evaluate topological stability. To target specific sites or loci that may be influencing relationships, we used site-wise and gene-wise likelihoods to identify which portions of the alignment drive topological differences among trees estimated with and without low coverage sites and with and without missing data. From these analyses, we produced a series of trees with different subsets of putative outliers removed and quantified the change in topology using a tree distance metric and support values, and summarized the information content of each locus. Next, we assessed whether the likelihood scores from the outlier analyses could be predicted by locus-specific alignment statistics. Finally, we took a more general data approach evaluating how data completeness impacted the estimated topology by producing a series of alignments with varying levels of missing data. The alternative data reduction approaches we employed allowed us to compare the utility of precise versus general filtering of missing data on phylogenetic inference. After rigorously assessing potential biases in the data we propose a phylogenetic hypothesis for lories and lorikeets.

## Materials and Methods

We sampled all 12 genera, 58/59 species, and 102/112 named taxa (Clements et al. 2019) within the Loriini, and three additional subspecies (*Glossopsitta concinna concinna*, *G. concinna didimus*, and *Trichoglossus haematodus caeruleiceps*) recognized by Gill and Donsker (2019) and Forshaw (2010), respectively. In total, we sampled 105 taxa within Loriini. *Charmosyna diadema* is the only species not included in our study, which is extinct and known from a single female specimen (Forshaw et al. 1989). Two additional taxa (*Eos histrio talautensis* and *E. squamata riciniata*) produced few loci with high missing data in those loci and were excluded from final analyses. We did not obtain samples from the following taxa: *C. rubronotata kordoana*, *Psitteuteles iris rubripileum*, *Neopsittacus pullicauda socialis*, *E. histrio challengeri*, *T. haematodus brooki*, and *T. moluccanus septentrionalis*. We treated *T. haematodus rosenbergii* as *T. rosenbergii* and *T. haematodus intermedius* as *T. haematodus haematodus* following Gill and Donsker (2019). We also followed Gill and Donsker (2019) and used *Parvipsitta* for *P. pusilla and P. porphyrocephala*. When possible, we sampled more than one individual per species to verify the phylogenetic position of a taxon. For outgroups, we used *Melopsittacus undulatus*, *Psittaculirostris edwardsii*, and *Cyclopsitta diophthalma*, which together with the Loriini form the clade Loriinae (Joseph et al. 2012; Provost et al. 2018). Sampling map and specimen details and locality information are available in supplementary fig. 1 and supplementary table S1.

We extracted total genomic DNA from muscle tissue using QIAamp DNeasy extraction kits (Qiagen, Valencia, CA). For historical samples from museum specimens, we used a modified DNeasy extraction protocol that used QIAquick PCR filter columns that size selected for smaller fragments of DNA. The modified protocol also included washing the sample with H_2_O and EtOH prior to extracting as well as extra time for digestion. DNA extraction from historical samples was done in a dedicated lab for working with degraded samples to reduce contamination risk. We quantified DNA extracts using a Qubit 2.0 Fluorometer (Thermo Fisher Scientific). Library preparation of UCEs and enrichment, and Illumina sequencing were performed by RAPiD Genomics (Gainesville, FL). The Tetrapod UCE 5K probe set was used to enrich 5,060 UCE loci (Faircloth et al. 2012). The wet lab component of this study was carried out over three years and the number of individuals multiplexed per lane ranged from 48–384. Sequencing was done on an Illumina HiSeq 2500 PE 125 or HiSeq 3000 PE 150. Fastq files are available on the Sequence Read Archive (SRA Bioproject ID:498485).

We used a modified data-processing pipeline that incorporated PHYLUCE (Faircloth 2015), a software package developed for analyzing UCE data, and seqcap_pop (Smith et al. 2014; Harvey et al. 2016). We used FastQ Screen to map raw reads to bacterial genomes and filter contaminant DNA (Wingett and Andrews 2018). Low-quality bases and adapter sequences were trimmed from multiplexed fastq files using Illumiprocessor v1 (Faircloth 2013; Bolger et al. 2014). Next, reads were assembled into contigs with Trinity v2.0.6 (Grabherr et al. 2011) and contigs were mapped to UCE probes. We chose the sample that produced the largest number of UCEs as the reference for subsequent mapping for all individuals. We generated an index of the reference sequence and independently mapped reads from each sample to the same reference sequence using BWA v0.7.13-r1126 (Li and Durbin 2009). SAM files produced from the BWA mapping were converted to BAM files, which were sorted with SAMtools (Li et al. 2009), and cleaned with Picard v1.106 (http://broadinstitute.github.io/picard). Then, we used the mpileup function in SAMtools to call variant sites and produce a VCF file, vcfutils to convert from VCF to fastq (excluding sites with quality scores < 20), and seqtk to convert fastq to fasta. From this last step, we produced two sets of DNA sequences that were analyzed independently. One dataset retained all variant sites irrespective of coverage (hereafter Low Coverage dataset), and a second set that excluded variant sites with less than 6x coverage using bcftools (hereafter Filtered dataset). These collective steps produced single fasta files containing all UCE loci for each individual sample. The following steps were independently performed for both datasets (Low Coverage and Filtered). Loci with greater than 30% missing sites were removed from each individual before alignment. In PHYLUCE we concatenated fasta files of each sample, aligned sequences in MAFFT (Katoh & Standley 2013), and retained loci where 75% of the samples were present in a locus for the final concatenated alignment.

### Low Coverage and Filtered outlier analysis

We estimated trees with 171 tips, which included multiple individuals per taxon. This dataset was used to check if samples from the same taxon grouped together as a means of identifying problematic samples. To focus on phylogenetic relationships among named taxa, all subsequent analyses are based on a reduced dataset that contained one sample per taxon with 105 ingroup taxa and three outgroup samples. We estimated phylogenomic trees for both the Low Coverage and Filtered concatenated alignments containing only unique taxa in IQ-TREE (Nguyen et al. 2014) using ModelFinder (Kalyaanamoorthy et al. 2017) to select the best-fit substitution model for each gene partition (Chernomor et al. 2016). To assess support, we estimated 1000 rapid bootstraps. The trees from the two different alignments did not produce the same phylogenetic relationships, so we performed an outlier sites/gene analysis to identify which sites were causing the topologies to be different. We performed a two-topology, site-specific log-likelihood test that estimated the site-likelihoods based on the gene partition of the Low Coverage alignment using topologies estimated from the Low Coverage (T_1_) and Filtered alignments (T_2_) in RAxML (Stamatakis 2014). We then estimated the change in site-wise log-likelihoods (hereafter Δ s-lk = T_1_ site log-likelihood - T_2_ site log-likelihood). We binned putative outlier sites into bins representing Δ s-lk: >20, >10, >2, <-2, <-10, and <-20. We produced new concatenated alignments that corresponded to each Δ s-lk threshold bin, for which outlier sites were converted to ambiguous sites (Ns) in all individuals. This approach allowed us to estimate trees with different levels of outlier sites removed from the alignment. Next, we converted the DNA sequence of each locus alignment into only parsimony informative sites using FASconCAT-G (Kück and Longo 2014), and summarized the amount of parsimony informative sites and missing data at these sites for modern and historical samples. To visualize how different the trees were, we measured the distance among 100 bootstrap trees using Robinson-Foulds distances (Robinson and Foulds 1981) with the multiRF function in phytools (Revell 2012) and used multidimensional scaling to plot the distances in two-dimensional space. All trees were processed and visualized using the R (R Core Team 2019) packages phytools and ape (Paradis et al. 2004). We classified samples into two categories: 1) samples that were collected within the last 30 years and came from frozen or ethanol preserved tissue (hereafter Modern), and 2) samples that came from dry museum skins with ages ranging from the 1960s through the late 1800s (hereafter Historical). To visualize the distribution of each sample type on the tree we colored tips blue (historical) or red (modern).

### Subclade outlier analyses

We performed a complementary outlier analysis assessing subclades, but in this set of analyses we compared trees estimated from alignments with and without missing data. By performing this analysis on subclades we were able to examine how missing data impacted different portions of the tree. This approach could not be applied to alignments containing all clades because at least one individual had missing data at every site in the alignment. The six clades including a single outgroup sample were based on preliminary phylogenetic analysis and were a) *Eos*, *Trichoglossus*, *Glossopsitta concinna*, and *Psitteuteles iris* (*n* = 22), b) *Parvipsitta* and *Psitteuteles* (*n* = 8), c) *Neospittacus* (*n* = 7), d) *Chalcopsitta* and *Pseudeos* (*n* = 14), e) *Lorius* (*n* = 18), and f) *Charmosyna*, *Vini*, and *Phigys* (*n* = 32). To produce a concatenated alignment for a clade, we followed the same steps listed above. To retain more sites in the larger clades we did not include redundant taxa or samples. We further reduced the sample size from 58 to 22 in the diverse clade containing *Eos*, *Trichoglossus*, *Psitteuteles iris*, and *Glossopsitta* because the amount of missing data in this clade was high. We estimated subclade trees in IQ-TREE following the same procedures described above to produce alternative topologies (T_1_ and T_2_) estimated from alignments with (T_1_) and without missing data (T_2_). Each tree was rooted with a single outgroup (*Oreopsittacus arfaki* for *Charmosyna*, *Vini*, and *Phigys*; *P. goldiei* for all other clades) using phyx (Brown et al. 2017). We performed the same site-specific log-likelihood procedure described above for two alternative topologies (T_1_ and T_2_) except that site-likelihoods were converted to gene-wise log-likelihoods to assess the impact of missing data across an entire locus by summing the site-likelihoods for each gene using the scripts in Walker et al. (2018). We then estimated the Δ gene-wise log-likelihoods (hereafter Δ g-lk).

To explore how these putatively biased loci impacted phylogenetic inference, we grouped loci into bins representing Δ g-lk scores of: > 2, > 10, > 20, < −2, and < −10 (there were no genes where Δ g-lk < −20 in the Filtered dataset). We followed the same procedure for producing concatenated alignments corresponding to each likelihood threshold bin, but for this step we excluded the outlier loci from the global concatenated alignment. We then estimated phylogenies from each alignment to assess how sensitive phylogenetic relationships were to excluding loci in each of these approaches. If missing data biases phylogenetic relationships then the exclusion of loci with positive Δ g-lk should alter relationships driven by missing data. The removal of loci with negative Δ g-lk will enhance relationships driven by biases in missing data.

### Trees with varying-levels of complete data

To determine at which percentage of data completeness was necessary to produce a topology similar to the Filtered tree, we generated a series of alignments from the Low Coverage dataset with increasing levels of complete data. Using trimal (Capella-Gutiérrez et al. 2009), we converted all missing sites to gaps (-) to conform to the software requirements and trimmed alignments by setting the percentage of individuals required to have an unambiguous site to retain the position in the alignment. In increments of 10%, we removed all sites where 0–100% of the sites had no missing data. This approach produced a range of alignments with no missing data (100%) through keeping all sites (0%). We estimated phylogenies for each of the eleven datasets in IQ-TREE using the same approach previously described except that we estimated best-fit model across the entire alignment because the gene partitions were not retained after filtering.

### Manipulating modern samples to mimic historical samples

To provide a complementary approach for assessing whether missing data versus data quality were biasing phylogeny we converted a percentage of sites in five modern samples (*Trichoglossus rubritorquis* KU22839, *T. chlorolepidotus* DOT2422, *T. ornatus* DOT7930, *Phigys solitarius* KU22543, and *Charmosyna placentis pallidior* DOT20055) to missing data. If the position of these samples was sensitive to the addition of missing data and the samples clustered with historical samples then missing data is the more likely culprit of the bias. We explored different percentages of missing data (50%–99.9%) and converting random sites versus only parsimony informative sites. We found that the position of these samples only changed when we converted 99.9% of parsimony informative sites to missing data, and we do not present results from the other thresholds. The reason why such a high percentage was necessary is because the position of these taxa were supported by a small number of parsimony informative sites, and the only way to influence the sites driving their relationships was to use a high threshold. We converted sites independently ten times and estimated phylogenies in IQ-TREE from the alignments using the same approach previously described.

### Summarizing phylogenetic signal in modern and historical samples

To explicitly compare the information content in DNA sequence from historical and modern samples, we calculated alignment statistics for each locus and for alignments of only parsimony informative sites partitioned into sample types using AMAS (Borowiec 2016). We compared the amount of parsimony informative sites and missing data in samples with their age. Because we sequenced samples over multiple years, batch effects, or biases attributable to differences among sequencing run could also bias our results (Leigh et al. 2018). To provide a qualitative assessment of batch effects, we provide plots of trees where tips have been colored according to one of three plates that they were sequenced on. If there were substantial batch effects in the data then phylogenetic relationships could be, in part, due to whether samples were sequenced together.

We built a neural net in the R package caret v. 6.0.79 (Kuhn 2008) to test whether the Δ gene-wise log-likelihood of each gene partition could be predicted by the alignment statistics. The alignment statistics (alignment length, number of undetermined characters, the number of parsimony informative sites, number of variable sites, and GC content) were specified as the input neurons, and the output neuron was the Δ log-likelihood. The input data were scaled to the min and max for each statistic, and the percentage of training/test data was set to 75%/25%, respectively. We produced 100 training/test datasets, independently ran each analysis, and report mean R^2^, RSME, and variable importance. We performed this analysis on the Low Coverage alignment that included all taxa and independently on the six sub-clades using the data from both filtering schemes.

## Results

### Data characteristics

We sequenced 176 unique samples, including 16 that were re-sequenced to improve the amount of data recovered. We dropped five individuals that had aberrant relationships and long branches in the tree, patterns that were presumably driven by limited data. The final dataset comprised 171 unique individuals (168 in the ingroup; three outgroups) where 54% and 46% modern were from historical and modern samples, respectively. Of the 58 species sampled, 27 had intraspecific sampling that included historical and modern samples. Historical samples (mean=5.5 million; SD=4.6 million), on average had more reads than modern samples (mean=3.53 million; SD=2.26 million), but a higher percentage of the reads in modern samples mapped to the reference (modern: mean=87.07%; SD =13.32%; historical: mean = 51.97 %; SD = 21.73%). In modern samples a greater number of sites were masked for having coverage < 6x (modern: mean=443,180; SD=202,764; historical: mean=359,958; SD=152,304). The mean per-site coverage across individuals was similar between the two sample types (modern: mean=67.85; SD=25.20; historical: mean=72.57; SD=28.15). Additional read and locus statistics are available in Supplementary Tables S2 and S3). We produced a Low Coverage dataset that included all variant sites irrespective of coverage, and Filtered dataset that excluded variant sites with less than 6x coverage. In the Low Coverage and Filtered datasets, loci had a mean length of 498 bps (range: 136–1,708 bps) and 482 bps (range: 105–1,413 bps), respectively. The mean and range number of taxa per locus was 164 for the Low Coverage (48–171) and 152 for the Filtered (5–171) datasets. After retaining loci where 75% of the individuals were present in any one locus, the Low Coverage dataset had 4,212 loci, 2,112,365 bps and 58,762 parsimony informative sites, whereas the Filtered concatenated alignment had 3,775 loci, 2,046,673 bps and 51,482 parsimony informative sites.

Overall, the Low Coverage dataset had more parsimony informative sites than the Filtered dataset (fig. 1). A comparison of sample types shows the range in number of parsimony informative sites among loci was lower in the modern samples in contrast to the historical samples (figs. 1A and 1B). In the Filtered dataset (fig. 1B) there was greater variability in the number of samples per locus in historical samples than the Low Coverage dataset (fig. 1A). For each alignment type, the modern samples contained a greater number of parsimony informative sites and less missing data than the historical samples (1.7x and 2.2x more parsimony informative sites in the Low Coverage and Filtered datasets, respectively; figs. 1C and 1D). In the Filtered dataset, the number of parsimony informative sites dropped, and the range of the number of individuals in each locus alignment increased. The percentage of missing data in samples decreased with specimen age (Adjusted R^2^ = 0.51; *n* = 144; p-value < 0.0001; supplementary fig. S2).

**FIG. 1.**
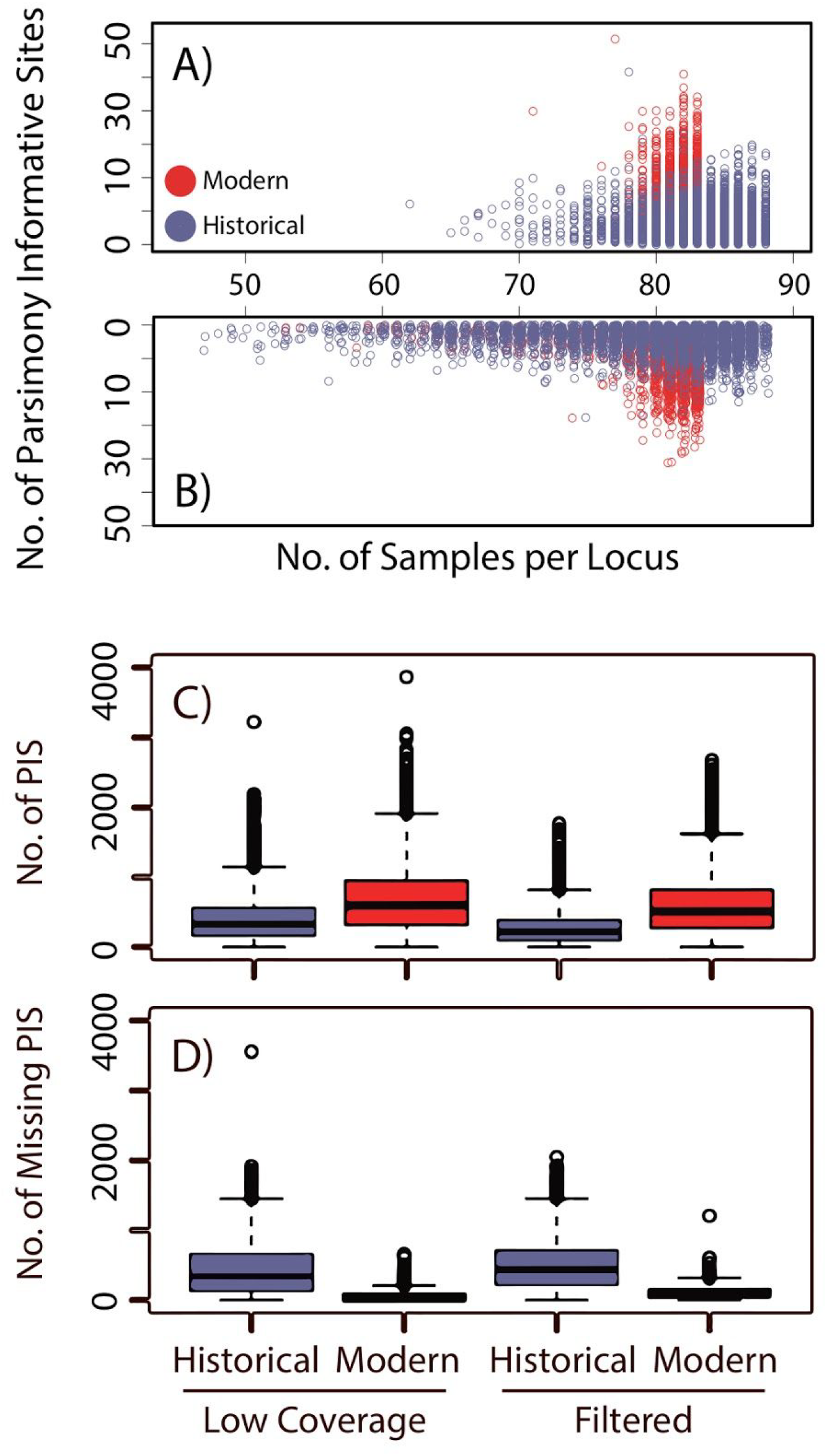
The information content varied among datasets and sample types. Shown is a Mirror plot with the number of samples and parsimony informative sites change between the Low Coverage (A) and Filtered (B) alignments. Each dot represents a locus and is colored according to whether the sample was from modern tissues (red) or historical specimens (blue). On the x-axis is the number of taxa in each locus alignment, and on the y-axis is the number of parsimony informative sites in each locus alignment. Panels C) and D) are boxplots showing the number of parsimony informative sites (PIS) and number of missing sites at parsimony informative sites, respectively.

### Resolving phylogenomic relationships among lorikeets

The backbone phylogeny we inferred for the Loriini generally had high support and the placement of genera was stable (supplementary figs. S3–S14). Summarizing higher-level relationships, *Oreopsittacus* was sister to all other ingroup taxa, then *Charmosyna* was sister to the clade containing *Neopsittacus*, *Lorius*, *Pseudeos*, *Chalcopsitta*, *Psitteuteles*, *Glossopsitta*, *Eos*, *Trichoglossus*, and *Parvipsitta*. The placements of *Neopsittacus*, *Lorius*, *Pseudeos*, and *Chalcopsitta* were well-supported in the tree, and each of these genera were monophyletic. *Trichoglossus*, *Charmosyna*, and *Psitteuteles* were not monophyletic. *Psitteuteles* was found in three separate places in the tree: *P. versicolor* was sister to the recently erected genus *Parvipsitta*; *P. iris* was nested within a clade of *Trichoglossus* taxa that are from Indonesia; and *P. goldei* was sister to the clade containing *Glossopsitta*, *Eos*, *Trichoglossus*, and *P. iris*. *Vini* and *Phigys* are strongly supported as nested within *Charmosyna*. Relationships within *Charmosyna* (including *Vini* and *Phigys*) and *Chalcopsitta* were generally stable across filtering schemes, as were relationships of the less diverse clades (*Oreopsittacus*, *Neopsittacus*, and *Parvipsitta*). Within the remaining clades, there were several notable differences in topological relationships among the Low coverage and Filtered trees.

The Filtered tree contains four clades containing *Trichoglossus*, *Eos*, *P. iris*, and *G. concinna* with varying levels of support (fig. 2A). *Glossopsitta concinna* was sister to a clade containing a monophyletic *Eos*, *Trichoglossus*, and *P. iris*. Within this tree, *Eos* was monophyletic and sister (bootstrap (BS) = 100%) to a clade containing *Trichoglossus* taxa that occur in Indonesia and the Philippines (*T. ornatus*, *T. flavoviridis*, and *T. johnstoniae*) and *P. iris*. The *Eos*, *Trichoglossus*, and *P. iris* clade was sister (BS = 85%) to a clade containing the remaining *Trichoglossus*, which was supported by a bootstrap value of 91%. This *Trichoglossus* clade had several short internodes and poorly supported relationships, particularly among *T. haematodus* subspecies, which primarily came from historical samples. *Trichoglossus euteles*, *T. forsteni*, *T. capistratus*, *T. weberi*, and *T. rubritorquis* are nested within *T. haematodus*. *Trichoglossus forsteni stresemanni* was more closely related to *T. capistratus* then to other *Trichoglossus forsteni* taxa. In contrast, the Low Coverage tree has two well-supported (BS > 95%) clades composed of *Trichoglossus*, *Eos*, *P. iris*, and *Glossopsitta concinna* (fig. 2B). One clade consists of entirely historical samples (*N* = 23), whereas the other was primarily modern samples (13/16). Within each of these clades, tips have similar relationships among taxa as seen in the Filtered tree. *Trichoglossus* that occur in Indonesia or the Philippines and *P. iris* are sister to *Eos* and the remaining *Trichoglossus* form a clade with the exception of one historical sample (*T. haematodus haematodus*). Support values are higher in the clade composed of mostly modern samples. In both trees (Low Coverage and Filtered), *Charmosyna* was composed of four clades, *C. wilhelminae* was sister to all other taxa in the clade, *C. rubronotata* and *C. placentis* are sister and form a clade, *C. multistriata* was sister to *C. josefinae* and *C. papou*, and the remaining *Charmosyna* taxa (*C. margarethae*, *C. rubrigularis*, *C. meeki*, *C. palmarum*, and *C. amabilis*) and *P. solitarius* and *Vini* form a clade. The position of *C. pulchella* and *C. toxopei* will be discussed below.

**FIG. 2.**
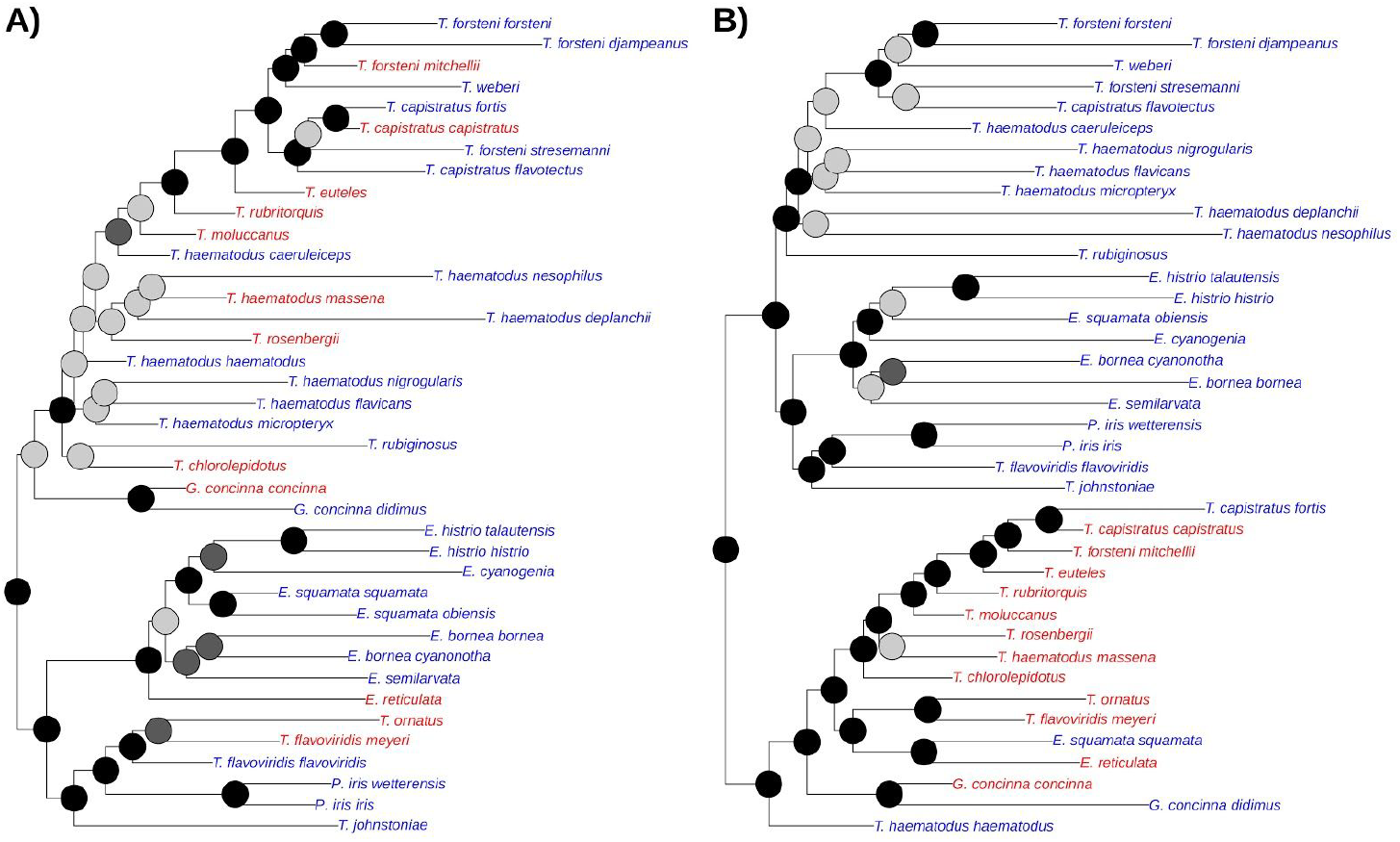
In the Low Coverage tree are clades composed of mostly historical versus modern samples. Shown is the subclade containing *Trichoglossus*/*Eos*/*Psitteuteles iris*/*Glossopsitta* from trees estimated without (A: Filtered Tree) and with low coverage sites (B: Low Coverage Tree). Bootstrap nodes are colored on a gradient from 100% (black) to < 70% (gray). Taxon names are colored according to whether their DNA came from modern tissues (red) or historical specimens (blue).

Similar clustering patterns based on sample type (historical versus modern) are observed in *Lorius*, *Vini*, and *Charmosyna* in the Low Coverage trees (supplementary fig. S3). The two subspecies of *L. lory* that come from modern samples are sister taxa. The one modern sample of *C. placentis* was sister to *C. rubronotata*. *Charmosyna palmarum* (a modern sample) was strongly supported as sister to *Phigys* and *Vini*. The three *Vini* from historical samples group together. The two *C. papou* subspecies that come from historical samples are sister to a clade containing the remaining *C. papou* subspecies. None of these relationships are observed in the Filtered tree.

A qualitative assessment of batch effects, by coloring each tip in the tree according to sequencing run, did not detect biases whereby samples would have clustered together based on sequencing plate (supplementary figs. S16 and S17). In the Low Coverage tree (supplementary fig. S18), biases in clustering were more apparent when tips are colored according to whether the sample came from a modern or historical source, which was not observed in the Filtered tree (supplementary fig. S19).

### Outlier sites and loci

The outlier analyses assessing the change in site-likelihoods scores between the Low Coverage versus the Filtered topology identified 3,142 (3,142 sites: Δ s-lk < −2; 489 sites: Δ s-lk < −10 = 489; 118 sites: Δ s-lk < −20) and 1,933 (1,933 sites: Δ s-lk > 2; 113 sites: Δ s-lk > 10; 4 sites Δ s-lk > 20) outlier sites in the alignment (2,112,365 bps) with negative and positive Δ s-lk values, respectively (fig. 3A). Higher and more positive Δ s-lk are sites that better support the topology estimated from the Filtered alignment, and lower and more negative values favor the tree estimated from the Low Coverage alignment. The 3,142 outlier sites with negative Δ s-lk were found on 1,904 loci, and the 1,933 outlier sites with positive values were on 1,378 loci. The 3,142 sites with Δ s-lk < −2, which favored the topology of the Low Coverage tree, exhibited a disproportionate number of missing sites in the historical versus modern samples (fig. 3B). We do not discuss the number of missing sites for sites Δ s-lk > 2 because positive sites favor relationships in the Filtered tree.

**FIG. 3.**
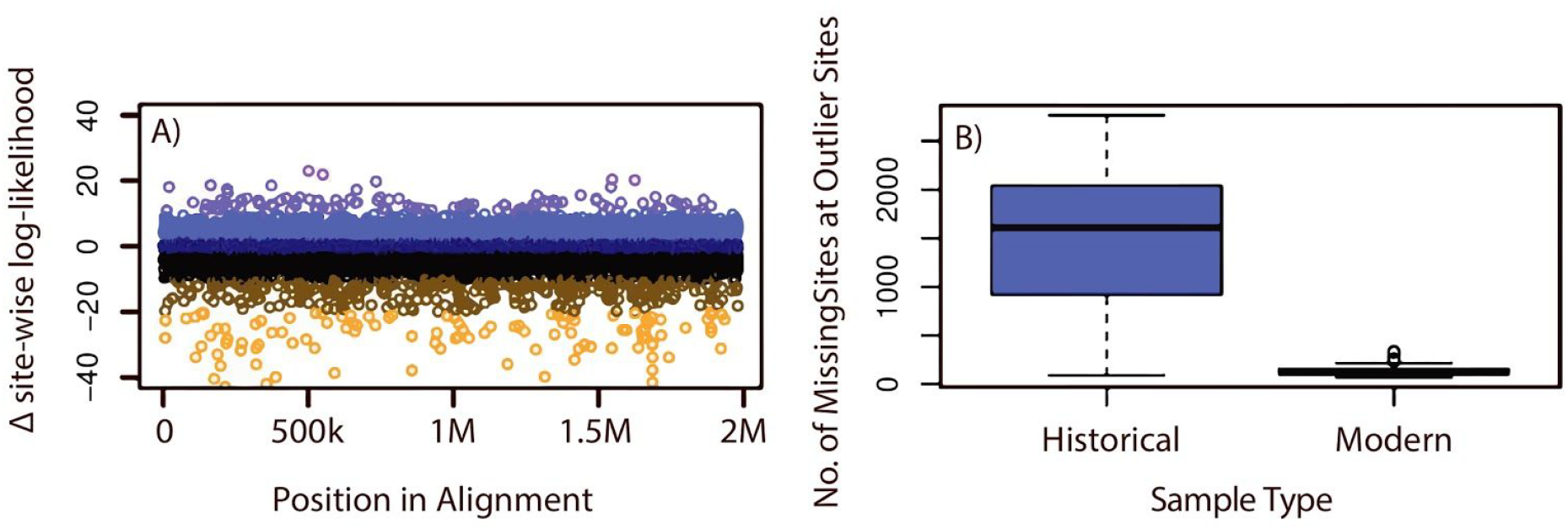
Outlier sites have high missing data in historical samples. A) Outlier site plot showing Δ sites-wise log-likelihoods (Δ s-lk) for topologies estimated with and without low coverage sites. The y-axis is the Δ s-lk score and the x-axis represents individual loci in the concatenated alignment, where k and M represent thousand and million, respectively. Points are colored according to the magnitude of the Δ site-wise log-likelihood scores according to a gradient ranging from > 20 (purple) through less than −10 (mustard). B) Boxplot of historical (blue) and modern (red) samples showing the amount of missing data in the 3,142 outlier sites (Δ s-lk < −2) identified in plot A.

Overall, the subclade outlier analyses for the Low Coverage alignment identified more outlier loci (fig. 2 and supplementary fig. S19). There were 47/4 (Low Coverage/Filtered), 371/35, and 1,730/518 loci in the three bins (Δ g-lk of: > 20, > 10, and > 2), respectively (fig. 4). There were 122/11 (Low Coverage/Filtered) and 1,336/316 loci in the three bins (Δ g-lk of: < −10 and < −2), respectively. The maximum and minimum Δ gene-wise log-likelihood were much higher in the Low Coverage (in *Trichoglossus*/*Eos*/*Psitteuteles*/*Glossopsitta*: Δ g-lk = −31.926–476.093) versus the Filtered dataset (in *Trichoglossus*/*Eos*/*Psitteuteles*/*Glossopsitta*: Δ g-lk = −18.247–41.402). There were 1,305 loci identified by both outlier analyses—the Low Coverage versus Filtered tree and subclade. In the Low Coverage and Filtered analysis, the outlier sites were found on 599 loci uniquely identified, and 425 loci identified by the subclade clade analyses.

**FIG. 4.**
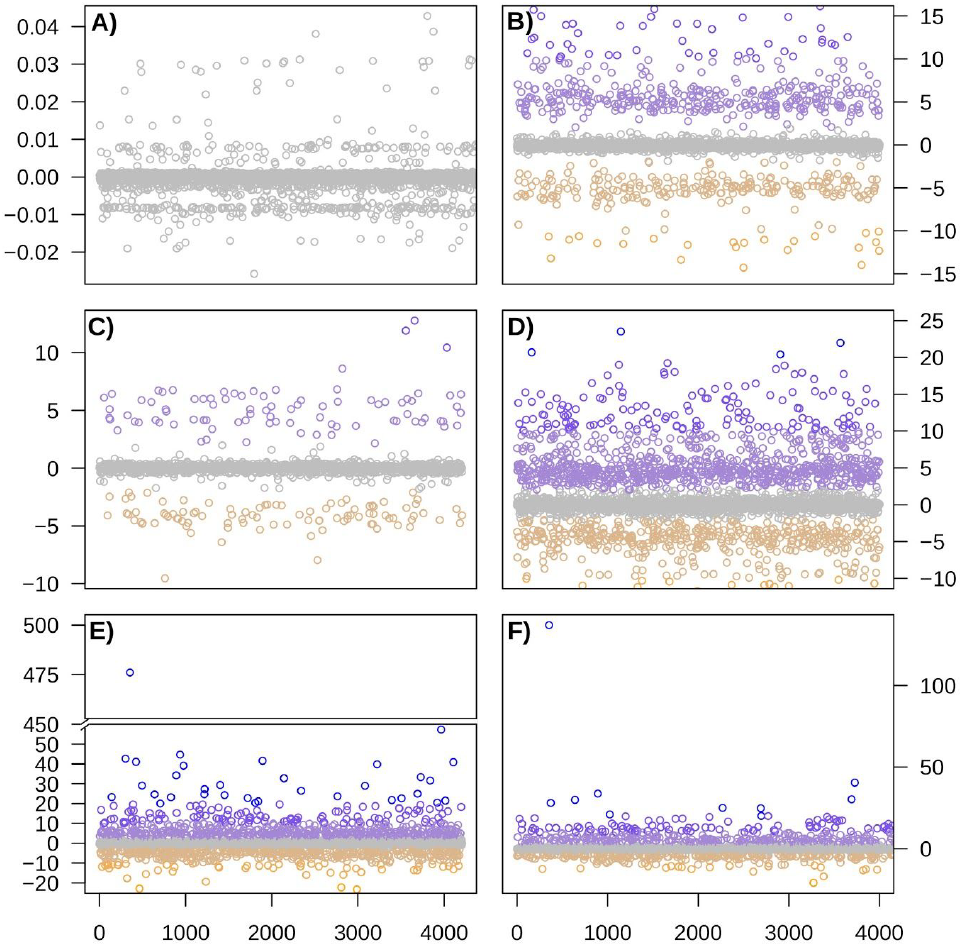
Likelihood plots showing Δ gene-wise log-likelihoods (Δ g-lk) for topologies estimated with and without missing data for the Low Coverage dataset. The y-axis is the Δ g-lk and the x-axis represents individual loci across the full alignment. Shown are the results for six subclades assessed within Loriini using the Low Coverage dataset: A) *Parvipsitta* and *Psitteuteles*, B) *Chalcopsitta* and *Pseudeos*, C) *Neospittacus*, D) *Charmosyna*, *Vini*, and *Phigys*, E) *Eos*, *Trichoglossus*, *Glossopsitta concinna*, and *Psitteuteles iris*, and F) *Lorius*. Points are colored according to the magnitude of the Δ g-lk scores according to a gradient ranging from > 20 (blue) through < −10 (orange).

We found that by converting parsimony sites to missing data, modern samples could cluster with historical samples. The extent of the shift of the sample and the position of the manipulated sample in the tree varied across the trees (supplementary figs S20A–H). For example, *T. ornatus*, which is strongly supported as sister to *T. flavoviridis* in the Low Coverage Tree (supplementary figs S3 and S20A), is nested within the clade containing only historical samples in the *Trichoglossus*/*Eos*/*Psitteuteles* clade in some of the trees with manipulated sequences (supplementary figs S20E, S20I, and S20J). *Charmosyna placentis pallidior* is strongly supported as sister to *C. rubronotata rubronotata*, and when its sites are converted to missing data it is nested within in its correct position in *C. placentis* (supplementary figs S20E, S20F, S20H, S20I, and S20K). In some trees, the position of taxa did not change at all (supplementary figs S20A-H), and in others the taxa were placed well outside their clade (supplementary figs S20A-H).

According to our neural net, alignment statistics were weak predictors of Δ g-lk scores in the Low Coverage versus Filtered trees (mean and SD R^2^ =0.05 (0.03); RMSE=0.03 (0.01) Δ g-lk scores. In the model GC content (Mean Variable Importance: GC content=31.54) and number of parsimony informative sites (25.14) were more important than the others (Alignment Length=12.89; No. of Taxa=12.89; Undetermined Characters=12.41; No. Variable sites=1.32). The subclade neural nets generally performed more poorly with R^2^ values ranging from 0.002–0.08, and parsimony informative sites and number of variable sites were the most important variables (Supplementary table S4). These results suggest that the loci with higher Δ g-lk values had higher parsimony informative sites, but there was no general trend between these variables across all loci.

### Impacts of filtering sites and loci

The removal of outlier loci with positive Δ g-lk scores broke-up some of the same-type clusters, particularly at a threshold value of all loci with Δ > 2 (supplementary figs. S4–S6). However, the removal of this many loci (*n* = 1,730) also reduced the support for other nodes in the tree. Trees estimated with the removal of negative outlier loci retained the apparent sample-type clusters (supplementary figs. S3, S7, S8). Individual taxa whose position varied the most among filtering schemes were *G. concinna* and *T. rubiginosus*. In the Filtered dataset, which did not exhibit the sample-type clusters, the removal of outlier loci (Δ g-lk > 2; supplementary fig. S12) increased the support for the placement of *G. concinna* as sister to the clade containing *Trichoglossus*, *Eos*, and *P iris*. In contrast, the removal of outlier loci (Δ g-lk < −2) placed *G. concinna* within the clade containing *Trichoglossus*, *Eos*, and *P iris*. This placement received moderate bootstrap support for either being sister to the clade containing *T. haematodus* and allies or the entire clade containing *Trichoglossus*, *Eos*, *P. iris*, and *G. concinna*. *Lorius lory* has seven subspecies, which formed a well-supported clade in the Filtered tree (supplementary fig. S9), with the exception of *L. lory viridicrissalis*, whose placement was equivocal. The Low Coverage tree has *L. lory viridicrissalis* within the *L. lory* clade with low support (supplementary fig. S3). Filtering of outlier loci changed support values, but never unequivocally placed *L. lory viridicrissalis* within *L. lory*. *Charmosyna pulchella* and *C. toxopei* are sister taxa, however, their position within *Charmosyna* varied across trees. Trees estimated with all loci or loci with negative Δ g-lk scores excluded had these taxa as sister (often with high support) to the clade containing the subclades *C. multistriata*; *C. josefinae* and *C. papou*; and *C. margarethae*, *C. rubrigularis*, *C. meeki*, *C. palmarum*, *C. amabilis*, *P. solitarius*, and *Vini* (supplementary figs. S7 and S8). Alternatively, trees where positive Δ g-lk scores were excluded had these taxa as sister, albeit with lower support (BS = 72% and 79%) to a clade containing *Charmosyna*, *Phigys*, and *Vini* (supplementary figs. S4–S6).

Examining the differences among topologies in multi-dimensional space showed distances among trees change across filtering schemes (fig. 5). In the trees where outlier sites were excluded (fig. 5A), the Robinson-Foulds distances among the Low Coverage and Filtered trees decreased. At a filtering threshold of Δ s-lk < −2 (3,142 sites), the distance between the two trees was minimal. Filtering sites with positive Δ s-lk values maintained a topology similar to the Low Coverage tree, except at a threshold of Δ s-lk > 2, which was distant from all other trees in multi-dimensional space. In the subclade analyses, the filtering of loci did not yield similar topologies between the Low Coverage and Filtered trees (fig. 5B). However, the Low Coverage tree where loci were excluded at a threshold of Δ s-lk > 2 was the least distant from the Filtered tree. The Low Coverage trees with all loci and Δ s-lk < −2 produced similar topologies and were the most distant from the Filtered trees. Despite some differences in the placement of taxa across the Filtered trees, the Robinson-Foulds distances among trees were comparatively low.

**FIG. 5.**
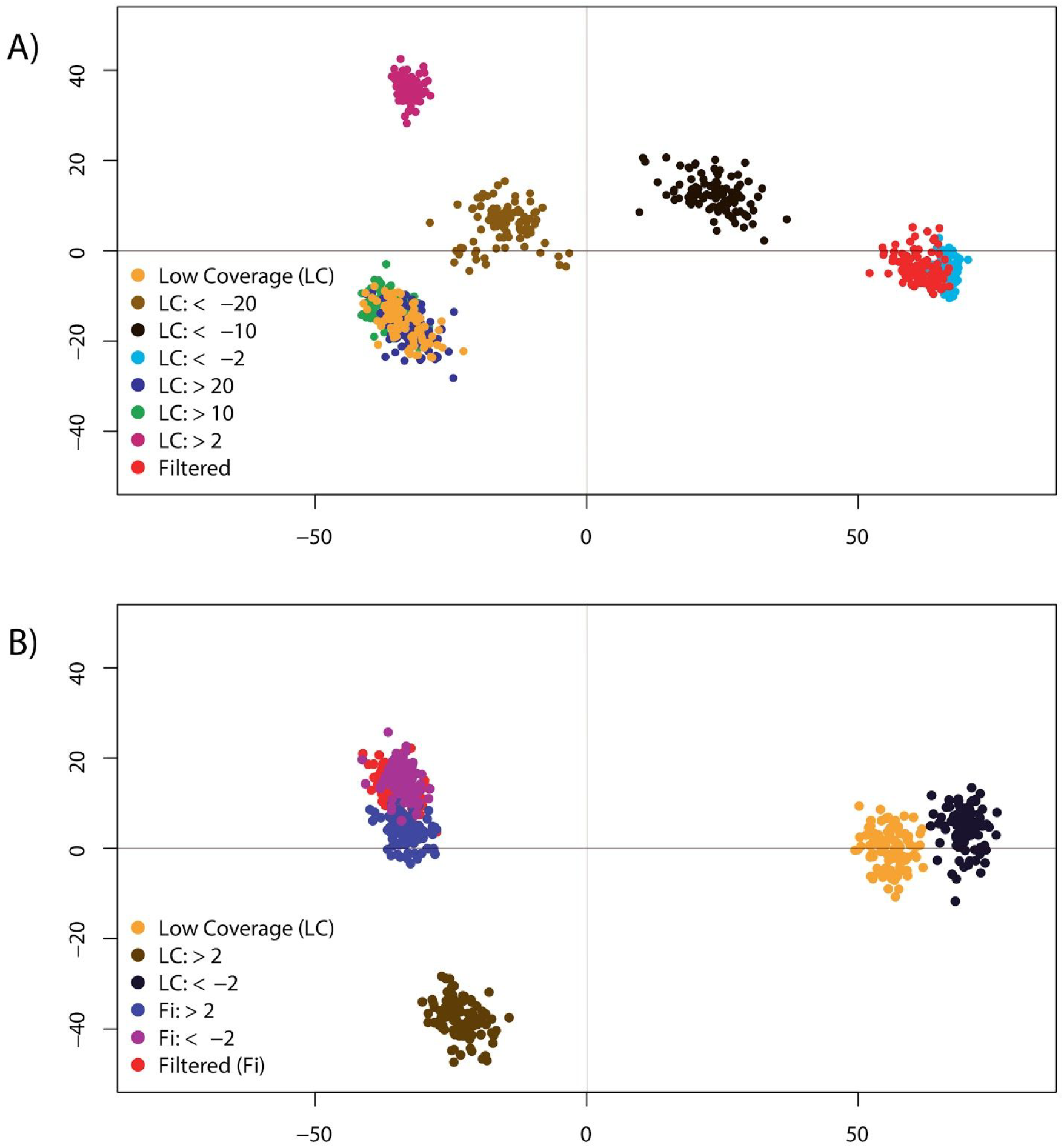
Multidimensional scaling of Robinson-Foulds distances among 100 bootstrap trees with differing levels of outlier sites or loci excluded. Plot A compares distances among Filtered and Low Coverage trees where outlier sites have been removed at different increments. Outlier sites were excluded in the Low Coverage alignment using Δ site-wise log-likelihood (Δ s-lk) thresholds of: >20, >10, >2, <-2, <-10, and <-20. Plot B shows the distances among trees produced from the subclade outlier analyses. Shown is a comparison of the Low Coverage and Filtered trees with topologies estimated with outlier loci excluded using Δ gene-wise log-likelihood (Δ g-lk) thresholds of: >2 and <-2.

### Impact of data completeness

The alignment length ranged from 2,106,278 bps (0% or all sites) through 41,504 bps (100% or no sites with missing data (Table 1)), and the trees estimated from these alignments are in supplementary figs. 21A–K. Across this same range of filtering, there were 30,380 (0%) through 373 (100%) parsimony informative sites (Table 1). At 60% completeness the sample-type clusters started to break-up (supplementary fig. S21G), and at 70% the tree was similar to the Filtered tree (supplementary figs. S9 and S21H). By 90% completeness some relationships differed from the Filtered tree (supplementary figs. S9 and S21J), and by 100% the tree had lower resolution and support (supplementary fig. S21K). Between the 70% to the 90% data completeness threshold the alignment was reduced from 1,186,092 bps to 1,024,262 bps, and 15,404 to 8,633 parsimony informative sites.

**Table 1.** Data completeness, alignment length, and number of parsimony informative sites at differing thresholds of missing data allowance. Shown are the percentage of individuals at each site with non-ambiguous characters across the Low Coverage alignment. The alignment length and number of parsimony informative sites (PIS) decrease the percentage of data completeness increases and more sites are excluded.

| **% of Data Completeness** | **Alignment Length (bps)** | **PIS** |
| --- | --- | --- |
| **0%** | 2,106,278 | 30,380 |
| **10%** | 1,901,690 | 30,160 |
| **20%** | 1,786,404 | 28,753 |
| **30%** | 1,744,411 | 27,934 |
| **40%** | 1,661,276 | 25,794 |
| **50%** | 1,498,375 | 21,763 |
| **60%** | 1,354,038 | 18,809 |
| **70%** | 1,186,092 | 15,404 |
| **80%** | 1,024,262 | 12,342 |
| **90%** | 800,132 | 8,633 |
| **100%** | 41,504 | 373 |

## Discussion

We showed that systematic bias caused by missing informative sites between DNA sequences from modern versus historical specimens can produce aberrant or unstable phylogenetic relationships. To obtain dense taxon sampling in our focal group, the Loriini, we leveraged samples collected over the last 100+ years and assessed how this sampling scheme impacted phylogenetic relationships by producing alignments with low coverage sites included and excluded. These two trees exhibited some striking differences. In the Low Coverage tree, there were numerous cases where historical or modern samples clustered together that were not observed in the Filtered tree (e.g., fig. 2). We employed a targeted and general approach to assess how missing data was influencing these unexpected relationships. The targeted method using a site outlier analysis showed that a small number of sites were driving the topological differences, and at these sites, historical samples had substantially more missing data (fig. 1). Excluding low coverage sites reduced the discrepancy in missing data between historical and modern samples, and at this level of disparity, the tree did not contain sample-type clusters and was similar in topology to the Filtered tree (fig. 5). A more nuanced look at outlier loci within subclades showed that many loci supported alternative topologies when sites with missing data were excluded, and the position of some branches shifted when these loci were excluded (supplementary figs. S3–S8). Biases in the historical samples could also be observed in modern samples by dropping the majority of their parsimony informative sites, which produced similar sample-type clusters observed in the Low Coverage tree (supplementary fig S20). Summary statistics of locus alignments were poor predictors of the likelihood scores used to identify outlier portions of the alignment, indicating that biased loci do not have characteristics differing from other loci (supplementary table 4). The general approach using the percentage of data completeness indicated that sites with high data completeness were necessary to avoid spurious relationships, but more stringent conditions of data completeness produced less resolved trees. After accounting for biased loci and understanding the stability of nodes, we inferred a more robust phylogenetic hypothesis for the Loriini. Taxonomic relationships within the clade can now be revised to reflect natural groupings, but for some groups, additional work is still necessary.

### Asymmetric information content among sample types

We found that alignments with high missing data produced biased phylogenetic relationships. In these trees, sample-type clades or sister relationships presumably formed because there was not enough information to place historical samples among the modern samples, but there was enough data to determine relationships among historical samples. Our analyses suggest that an asymmetry in phylogenetic information content among sample types is the primary culprit of the bias because only 3,142 sites (6.6% of total sites) drove the topological differences among trees, and the historical samples had 7.5x more missing data at these sites (fig. 3). By filtering for data completeness, we produced a similar result and inferred the expected phylogeny by including only sites where 70% of the individuals had unambiguous characters. Previous work has shown that ambiguous sites can bias the probability of taxa being sister (Lemmon et al. 2009) and increase the resolution and support of clades (Simmons 2012; Simmons 2014). These previous studies did not deal with historical versus modern samples, and did not have the magnitude of characters in our dataset, but a similar mechanism is likely operating. Although, we accounted for among-site rate variation, which has been shown to lead to biases in missing sites (Lemmon et al. 2009), we did not evaluate how multispecies coalescent approaches would deal with our dataset. We concentrated instead on a concatenated approach, because our data met two criteria in which species-tree summary methods perform poorly (Molloy and Warnow 2018); namely, our data comprised 1) many poorly resolved gene trees with high missing data from 2) loci with low information content (UCEs).

By including low coverage sites we were able to explore potential biases that can arise between historical and modern samples. Filtering according to a read coverage threshold at each variant site is common practice in population genomic studies (e.g., Thom et al. 2018), but this approach is less frequently employed in phylogenomic bioinformatic pipelines (e.g., Faircloth 2015). In the Low Coverage tree, we found clusters of historical or modern samples that were not present in the Filtered tree (fig. 2; supplementary figs S3 and S9). Besides an asymmetry in informative sites, these clusters could be caused by sequencing errors present in one sample type, batch effects, or contamination. We address sample type in detail below, but we suspect that biases of batch effects and contamination were minimal. For example, we had limited power to test for batch effects because we did not randomly and evenly sequence samples across runs, therefore, there are portions of the tree where clades are composed almost entirely of samples from the same sequencing run (supplementary figs. S15 and S16). In these cases, we do not interpret these patterns as batch effects because the tips occur in their expected topological position and samples from different sequencing lanes are distributed throughout the tree. We took great care to avoid contamination during wet-lab procedures (Mundy et al. 1997) and we have no strong reason to suggest that contamination is driving the observed pattern, particularly after exploring the impacts of missing data on the topology. The impact of contamination may have been more pronounced on low quality sites, which were filtered out in all treatments, because unreported preliminary trees estimated with these low quality sites produced trees with long branches. However, more subtle effects of contamination on a small number of sites may not be directly detectable in the approaches we employed. Although we cannot rule out additional artifacts caused by contamination or sequencing error, the topology within each of the most apparent sample-type clusters in the *Trichoglossus*/*Eos*/*Psitteuteles* clade exhibited the expected relationships among taxa.

Our outlier analysis on subclades also found loci that were impacted by missing data. In the Low Coverage tree, the sample-type clusters were broken up when outlier loci with positive values were excluded but also reduced support values (supplementary figs. S3–S8). The exclusion of outlier loci with negative values retained the biased relationships. These loci had negative values because the topologies estimated with all sites with missing data removed were more likely given the alignment. The removal of these loci produced alignments that only included loci that were either biased or not impacted by missing data. In the Filtered dataset, the number of identified outlier genes was reduced and exclusion of outlier loci was less profound. Nonetheless, the removal of outlier loci in the Filtered dataset showed how the placement of *G. conccina*, *T. rubiginosus*, and the clade containing *C. pulchella* and *C. toxopei* was sensitive to missing data (supplementary figs. S9–S14). Interestingly, about 75% of the loci identified by the subclade outlier analyses were the same loci of the outlier sites identified by the Low Coverage versus Filtered outlier analysis. The information content of the non-overlapping loci is important because the more targeted site-wise outlier analysis was better at reconciling topological differences among the Low Coverage and Filtered trees than was the subclade approach.

There was a tendency for historical samples to fall outside of their clade or even the ingroup, as evident in previous phylogenomic studies on birds (Hosner et al. 2016; Moyle et al. 2016; Andersen et al 2019; McCullough et al. 2019). This was the case for seven of our excluded samples, which produced limited data and could not be accurately placed in their genus or higher-level clade. The sample-type clusters within *Trichoglossus*/*Eos*/*Psitteuteles* is an extreme example of this pattern, and the pattern is so striking because of the high number of historical samples in this particular clade (fig. 2). In the trees wherein a subset of modern samples we converted most parsimony informative sites to missing data, we observed the same pattern whereby some of the manipulated samples were inferred outside of their expected clades (e.g., supplementary fig. S20J). Without prior information on whether a taxon is sister or falls outside of a clade of closely related taxa, samples with high missing data in large alignments may not be able to be accurately placed on a phylogeny.

### Identifying biased samples and loci

Our neural net models were not able to predict Δ g-lk scores, but the number of parsimony informative sites was often the most important variable in the models (supplementary table S4). An explanation for the low predictability of the models was that even though the loci with high Δ g-lk scores tended to have a higher number of parsimony informative sites, there was also a large number of loci with high information content with low scores. The collective findings of the neural net models suggest that systematic bias associated with the characteristics of a locus alignment such as GC content or sequence length did not explain why loci were detected in the outlier analyses. Instead, the inability of the model to predict outlier scores indicates that biases in the loci were randomly distributed across phylogenetically informative loci. Because the magnitude of the Δ g-lk score is going to be in part dictated by how much information is at a site or across a locus, the outlier analysis is expected to identify sites or loci that have enough information to distinguish alternative trees. However, because missing data at less informative sites is also known to bias phylogenetic inference (Simmons 2012; Simmons 2014), the outlier analysis we used may not capture the full extent of missing data on our inferred phylogenies.

Preferentially selecting phylogenetically informative loci is expected to produce trees with better support (Gilbert et al. 2018), but our results suggest that this practice can produce less reliable relationships when the data content dramatically varies among samples. Other work has shown that filtering phylogenomic markers by information content had mixed results in terms of resolving discordance among trees estimated with different phylogenetic methods (Mclean et al. 2018). Outlier analysis using site and gene likelihood scores (Shen et al. 2017; Walker et al. 2018) provide a rapid means of identifying loci that have a large impact on phylogeny reconstruction, but, as we showed, the resolution of this approach will depend on the trees that are available for comparison (e.g., the a priori expected phylogeny versus an alternative phylogeny). As mentioned above, a targeted outlier approach will not address all potential biases that missing data can cause, but it can identify sites that are having a strong influence on the phylogeny. Despite the limitations of site-likelihoods, the precision of identifying specific sites/loci may be the more favorable option to filtering data because the alternative of using percentage of data completeness to remove sites resulted in removing sites that were important for other portions of the tree. This idiosyncratic behavior of filtering for data completeness to achieve higher topological support for one recalcitrant historical sample occurred in a recent study of honeyeaters. Andersen et al. (2019) increased the filtering stringency towards more complete datasets to improve support for *Gymnomyza aubryana*, however, previously well-supported nodes elsewhere in the tree were negatively impacted due to a reduction in total parsimony informative sites. The optimal percentage of data completeness will vary among datasets and depend on how asymmetric the information content is among sample types. For our dataset, there was a narrow window for when data completeness produced a reliable phylogeny (e.g., 70% versus 90%; supplementary figs S21H and S21J) because data completeness > 70% led to a less-resolved tree.

### Taxonomic implications

Our study builds on previous phylogenetic work on the Loriini by further clarifying relationships and adding 64 previously unsampled taxa (fig. 6). We inferred a backbone phylogeny of relationships among genera that was fairly well-resolved with the exception of the clade containing *Trichoglossus*, *Psitteuteles iris*, *Eos*, and *Glossopsitta*, and some nodes in *Charmosyna*. Our analyses corroborated recently proposed taxonomic changes where *Pseudeos cardinalis* was moved into *Pseudeos* from *Chalcopsitta*, and *Parvipsitta* was resurrected to contain *P. pusilla* and *P. porphyrocephala*, which were previously placed in *Glossopsitta* (Schweizer et al. 2015). In all of our trees, *P. fuscata* and *P. cardinalis* were sisters and were in turn sister to *Chalcopsitta*. *Parvipsitta pusilla* and *P. porphyrocephala* were sisters and not closely related to *G. concinna*. However, we found strong support for *P. pusilla* and *P. porphyrocephala* being sister to *Psitteuteles versicolor*, a novel result. *Psitteuteles versicolor* and *Parvipsitta* could be subsumed under a single genus. Irrespective of this taxonomic decision, the polyphyly of *Psitteuteles* will require that *P. goldei* and *P. iris* be moved into new genera. *Psitteuteles goldei* is sister to the clade containing *Trichoglossus*, *Eos*, *P. iris*, and *Glossopsitta*. The taxonomic revision of *P. iris* will depend on how *Trichoglossus* is treated, because *P. iris* is nested within a geographically coherent clade of taxa distributed largely to the west of New Guinea. The clade containing *Charmosyna*, *Phigys*, and *Vini* represents a deep, diverse, and geographically widespread group. The species in these genera are varied in terms of body size and shape, tail length, plumage color, and sexual dimorphism (Forshaw et al. 1989; Merwin et al. 2020), and these morphological traits are not found in monophyletic groups in our phylogeny. Species-level relationships among species in *Charmosyna* were well-supported and stable with the exception of the placement of *C. toxopei* and *C. pulchella*. Overall, the taxonomic revision of this clade will present challenges regarding when and where to split or lump taxa and how best to circumscribe genera.

**FIG. 6.**
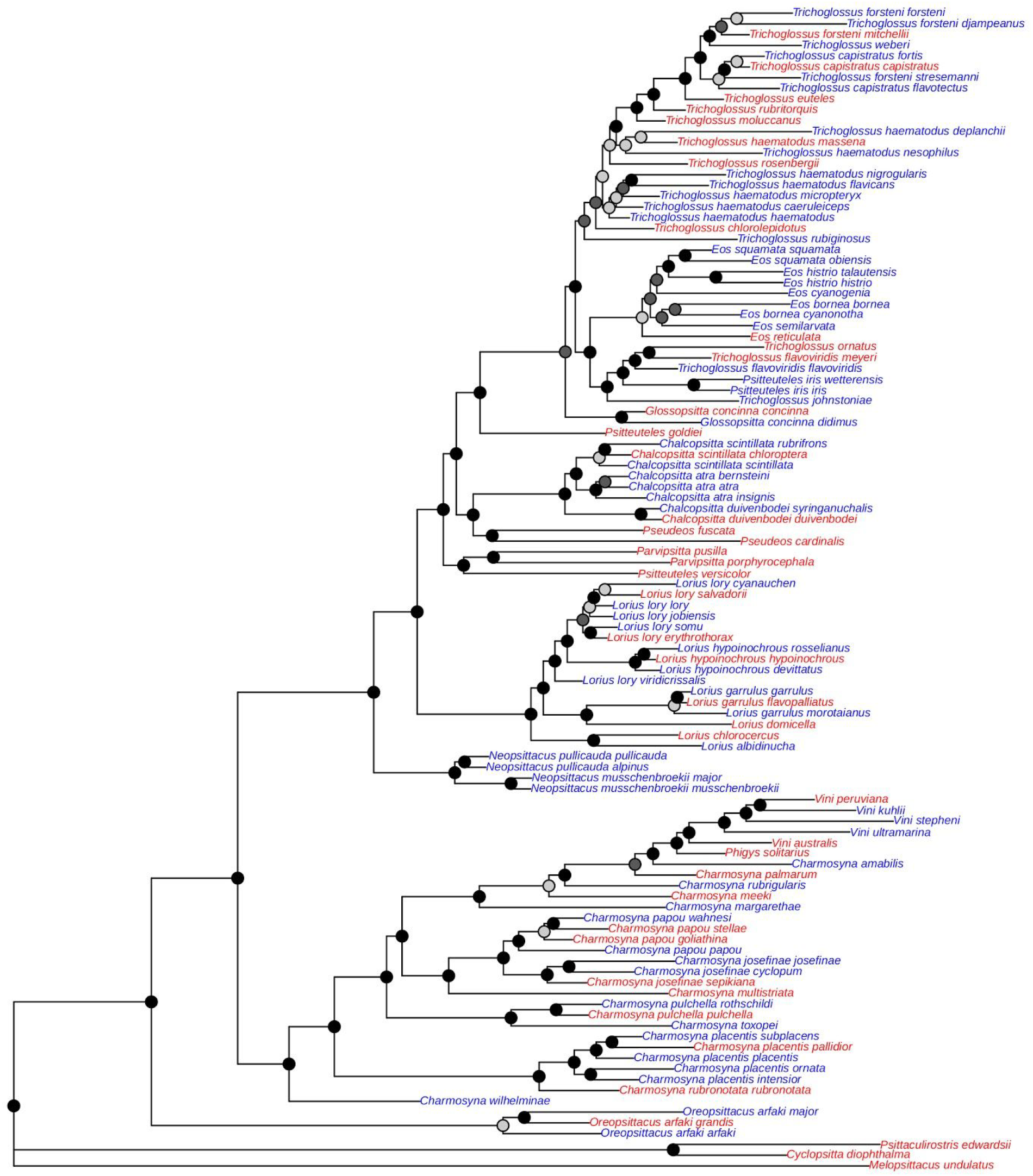
Maximum likelihood tree containing unique taxa in Loriini. The tree was inferred from a concatenated alignment where loci identified with the gene likelihood analysis with Δ gene-wise log-likelihood (Δ g-lk) values of > 10 were excluded. On each node are shown rapid bootstrap values and the taxon names are colored according to whether their DNA came from modern tissues (red) or historical specimens (blue). Bootstrap nodes are colored on a gradient from 100% (black) to < 70% (gray).

Relationships among subspecies within species varied substantially among taxa. Support for relationships among species within *Lorius* were generally stable except the placement of *L. lory viridicrissalis*, which was only nested within *L. lory* in the Low Coverage tree. Our *L. lory viridicrissalis* was a historical sample with a high degree of missing parsimony informative sites and its position as sister to *L. lory* and *L. hypoinochrous* is most likely an artifact. There were also varying levels of support for relationships among the other subspecies in *L. lory*, the most diverse species in the genus. Relationships among subspecies in *C. papou*, *C. josefinae*, and *C. placentis* had high support. Our analyses inferred a paraphyletic *T. haematodus* (with low support) and *T. forsteni*, the latter of which is still included in *T. haematodus* by some taxonomic checklists (Clements et al. 2019; Dickinson and Remsen 2013). This clade had many historical samples, which likely contributed to the clade’s low support, but even several of the taxa from modern samples were not placed with high support in the clade. Resolving these challenging relationships within *Trichoglossus* will likely require finer-resolution genetic data and expanded population-level sampling.

## Conclusion

Next-generation sequencing has provided systematists with an unprecedented amount of information for inferring phylogenetic relationships (McCormack et al. 2013). However, phylogenomic datasets are being produced faster than the development of best practices for assembling, processing, and analyzing large datasets for phylogenetic inference, particularly as the use of low-quality museum samples increases. Alignments produced without careful inspection may harbor biased loci that can have a large impact on downstream analyses (Springer and Gatesy 2018). Our findings have general implications for phylogenomic studies where there is an asymmetry in parsimony informative sites among closely related taxa. Although missing data have shown ambiguous impacts on phylogenetic inference (Lemmon et al. 2009; Simmons 2012; Wiens and Morrill 2011; Hovmöller et al. 2013; Simmons 2014; Streicher et al. 2015), the combination of a much higher number of informative sites in contemporary phylogenomics with an asymmetry between samples of different quality warrants new investigations on biases that can arise in alignments. The magnitude of biases will likely vary according to clade diversity and age, and the number of loci collected. We found that the bias was most extreme in a diverse and rapid radiation where there was likely limited information, even in complete loci, for teasing apart relationships. Shallow systematic and phylogeographic studies are expected to be the most difficult temporal scale for resolving relationships when there are high missing data associated with particular samples. Moving forward, having an understanding of the informational content of a locus, and how that information affects genealogy, will help avoid inferring dubious phylogenomic relationships.

## Acknowledgments

We thank the following institutions and people for providing the material used in this study: AMNH (P. Sweet, T. Trombone, G. Rosen, A. Caragiulo), UWBM (S. Birks, R. Faucett, J. Klicka), USNM (B. Schmidt, H. James, G. Graves), ANWC (R. Palmer, L. Joseph), LSUMZ (D. Dittmann, S. Cardiff, R. Brumfield, F. Sheldon), FMNH (B. Marks, J. Bates, S. Hackett), KU (M. Robbins, R. Moyle). This study was funded by National Science Foundation awards to BTS (DEB-1655736) and MJA (DEB-1557051). We also thank F. Burbrink, K. L. Provost, and J. Merwin for providing code, and J. Merwin, K. L. Provost, L. R. Moreira, G. Thom, B. Faircloth, C. Oliveros, M. Harvey, and anonymous reviewers for feedback and suggestions.

## SUPPLEMENTARY TABLE LEGENDS

**Supplementary Table S1. Metadata for Loriini samples used in this study.**

**Supplementary Table S2. Read and locus statistics.**

**Supplementary Table S3. Mean coverage of each site along a locus.**

**Supplementary Table S4. Neural Net model output for subclades that assessed predictors of Δ gene-wise log-likelihood scores** Shown are the variable importance of the five alignment statistics included in the neural net model for each subclade in Loriini. a) *Eos*, *Trichoglossus*, *Glossopsitta concinna*, and *Psitteuteles iris*, b) *Parvipsitta* and *Psitteuteles*, c) *Neospittacus*, d) *Chalcopsitta* and *Pseudeos*, e) *Lorius*, and f) *Charmosyna*, *Vini*, and *Phigys*. The per locus statistics were as follows: parsimony informative sites (PIS), alignment length (Ali. Len.), number of variables sites (Var. Sites), number of undetermined characters (Undet. Char.), and GC content (GC. content). The sample size for each subclade was the total number of alignments and 75%/25% was used to train/test each model. Reported are R^2^ and mean square error (MSE).

## SUPPLEMENTARY FIGURE LEGENDS

**Supplementary Figure S1.** Sampling map of lory and lorikeet taxa used in this study. Colored symbols represent material that came from historical (blue) and modern (red) samples.

**Supplementary Figure S2. Plot showing that older samples have more missing data.** Shown is the year the sample was collected at the percentage of missing characters as determined by IQ-TREE. Mean regression line and 95% CI are shown (Adjusted R^2^ = 0.51; *n* = 144; p-value < 0.0001). Not all samples were included in this analysis because accurate ages of the specimens were unavailable.

**Supplementary Figure S3–S8. Low Coverage maximum likelihood trees.** The tree was inferred from a concatenated alignment with all loci (fig. S3) and an increasing number of loci excluded in increments of the Δ gene-wise log-likelihood scores of: (Δ s-lk) thresholds of: >20 (fig. S4), >10 (fig. S5), >2 (fig. S6), <-10 (fig. S7), and <-2 (fig. S8). On each node are shown bootstrap values. Tip labels are colored according to whether the sample came from a historical (blue) or modern (red) sample.

**Supplementary Figure S9–S14. Filtered maximum likelihood trees.** The tree was inferred from a concatenated alignment with all loci (fig. S9) and an increasing number of loci excluded in increments of the Δ gene-wise log-likelihood scores of: (Δ s–lk) thresholds of: >20 (fig. S10), >10 (fig. S11), >2 (fig. S12), <-10 (fig. S13), and <-2 (fig. S14). Bootstrap values are shown on each node. Tip labels are colored according to whether the sample came from a historical (blue) or modern (red) sample.

**Supplementary Figure S15 and S16. Low Coverage and Filtered maximum likelihood trees with all samples (*N* = 171) and tips colored by sequencing plate.** Presented are Low Coverage (fig. S15) and Filtered (fig. S16) trees where tips were colored according to one of three (Plate 1: red; Plate 2: orange; and Plate 3: blue) lanes, which were sequenced independently over the course of three years.

**Supplementary Figure S17 and S18. Low Coverage and Filtered maximum likelihood trees with all samples (*N* = 171) and tips colored according to sample type.** Presented are Low Coverage (fig. S17) and Filtered (fig. S18) trees where tips are colored according to whether the sample type came from a historical specimen (blue) or modern tissue (red).

**Supplementary Figure S19. Likelihood plots showing gene Δ gene-wise log-likelihoods for topologies estimated with and without missing data for the Filtered dataset.** The y-axis is the Δ gene-wise log-likelihood and the x-axis represents individual loci across the full alignment. Shown are the results for six subclades assessed within Loriini using the Filtered dataset: A) *Parvipsitta* and *Psitteuteles*, B) *Chalcopsitta* and *Pseudeos*, C) *Neospittacus*, D) *Charmosyna*, *Vini*, and *Phigys*, E) *Eos*, *Trichoglossus*, *Glossopsitta concinna*, and *Psitteuteles iris*, and F) *Lorius*. Points are colored according to the magnitude of the Δ gene-wise log-likelihood scores using a gradient ranging from > 20 (blue) through < −10 (orange).

**Supplementary Figure S20. Low Coverage maximum likelihood trees where 99.9% of parsimony informative sites in five modern individuals were converted to missing data.** Shown are 11 trees where the sequence was not manipulated in tree A and trees B–K were. Tips are colored according to whether the sample type came from a historical specimen (blue) or modern tissue (red), and black samples are the ones that were manipulated (*Trichoglossus rubritorquis* KU22839, *T. chlorolepidotus* DOT2422, *T. ornatus* DOT7930, *Phigys solitarius* KU22543, and *Charmosyna placentis pallidior* DOT20055). On each node are shown rapid bootstrap values.

**Supplementary Figure S21. Low Coverage maximum likelihood trees with varying levels of data completeness.** Shown are 11 trees (A–K) estimated from alignments where the percentage of data completeness ranged from 0% (tree A; all sites were used) through 100% (tree K; all sites with missing data were excluded). In increments of 10%, each successive tree had higher data completeness. At each site along the alignment a specified percentage of individuals were required to have the site in order to be retained in the alignment. The trees are as follows and the percentages refer to the level of data completeness–A:0%; B:10%; C:20%; D:30%; E:40%; F:50%; G:60%; H:70%; I:80%; J:90%; and K:100%. Tips are colored according to whether the sample type came from a historical specimen (blue) or modern tissue (red),

